# Beta-band desynchronization reflects uncertainty in effector selection during motor planning

**DOI:** 10.1101/2021.04.23.441147

**Authors:** Milou J.L. van Helvert, Leonie Oostwoud Wijdenes, Linda Geerligs, W. Pieter Medendorp

## Abstract

While beta-band activity during motor planning is known to be modulated by uncertainty about where to act, less is known about its modulations to uncertainty about how to act. To investigate this issue, we recorded oscillatory brain activity with EEG while human participants (n = 17) performed a hand choice reaching task. The reaching hand was either predetermined or of participants’ choice, and the target was close to one of the two hands or at about equal distance from both. To measure neural activity in a motion-artifact-free time window, the location of the upcoming target was cued 1000-1500 ms before the presentation of the target, whereby the cue was valid in 50% of trials. As evidence for motor planning during the cueing phase, behavioral observations showed that the cue affected later hand choice. Furthermore, reaction times were longer in the choice than in the predetermined trials, supporting the notion of a competitive process for hand selection. Modulations of beta-band power over central cortical regions, but not alpha-band or theta-band power, were in line with these observations. During the cueing period, reaches in predetermined trials were preceded by larger decreases in beta-band power than reaches in choice trials. Cue direction did not affect reaction times or beta-band power, which may be due to the cue being invalid in 50% of trials, retaining effector uncertainty during motor planning. Our findings suggest that effector uncertainty, similar to target uncertainty, selectively modulates beta-band power during motor planning.

**New & Noteworthy:** While reach-related beta-band power in central cortical areas is known to modulate with the number of potential targets, here we show, using a cueing paradigm, that the power in this frequency band, but not in the alpha or theta-band, is also modulated by the uncertainty of which hand to use. This finding supports the notion that multiple possible effector-specific actions can be specified in parallel up to the level of motor preparation.

## Introduction

At a picnic with many delicacies, there are numerous opportunities for action. We can look at one of several treats, or reach for it, and when we reach, we could use the left or right hand. How is this decision process being solved? Computational theories suggest that the brain chooses the action that maximizes utility, which depends on the cost associated with performing the action and the desirability of the outcome, i.e., the reward (Haggard, 2008; Shadmehr, Huang, & Ahmed, 2016; Wolpert & Landy, 2012). In neural terms, it follows that the circuits involved in deciding between actions based on utility are strongly coupled to the circuits responsible for generating an action. Indeed, neurophysiological studies have suggested that multiple potential motor plans can be encoded in parallel and compete for selection within the brain’s sensorimotor regions (Cisek, 2006).

In non-human primates, most of the evidence for this process of embodied decision making comes from experiments that manipulated the number or location of potential targets (Basso & Wurtz, 1997; Cisek & Kalaska, 2005; Glaser, Perich, Ramkumar, Miller, & Kording, 2018; Klaes, Westendorff, Chakrabarti, & Gail, 2011). For example, in a unimanual reaching task with two potential targets, neural activity in dorsal premotor cortex represents both options simultaneously and reflects the selection of one over the other when the choice is made (Cisek & Kalaska, 2005; but see Dekleva, Kording, & Miller, 2018 for an alternative interpretation). Analogous results have also been observed in humans. For instance, Tzagarakis et al. (2010, 2015) reported that cortical beta-band desynchronization, associated with motor planning (Jasper & Penfield, 1949; Pfurtscheller, 1992), depends on the number of potential targets and their directional uncertainty. Grent-’t-Jong et al. (2014; 2015) reported that the proximity of two potential reach goals has a direct influence on motor cortex activity, as measured by oscillatory power (see also Tzagarakis et al., 2015).

Utility of a movement does not only depend on the location of the target, it is also determined by the effector that needs to be moved. Within this notion, target and effector selection can be considered as part of an integrated computation in movement planning, in which the expected utility of each potential movement is defined by the distance and direction of the respective target relative to the respective effector (Bakker, Selen, & Medendorp, 2018; Dancause & Schieber, 2010; Schweighofer et al., 2015). Accordingly, if multiple potential targets evoke multiple concurrent movement plans of a single effector, deciding between multiple effectors to move to a single target may also lead to the specification of parallel movement plans. This has been indeed observed when selecting between eye versus arm movements; cortical areas involved in these movements are simultaneously activated until the effector is selected, as observed both in monkeys (Cui & Andersen, 2011) and humans (Medendorp & Heed, 2019, for review). However, it is important to realize that eye and hand movements serve different purposes and, in natural situations, are typically used in combination (Heed, Beurze, Toni, Röder, & Medendorp, 2011), which could explain their simultaneous specification.

It is less clear whether the brain simultaneously specifies motor plans for the two arms. Using a combined EEG-fMRI study, Bernier et al. (2012) tested participants in an arm choice experiment with a fixed target location, and found activity in parietal and premotor cortex only contralateral to the reaching arm after target onset. This could be interpreted as if effector selection precedes movement planning, i.e. that hand selection is not associated with the simultaneous specification of two motor plans. This would be in line with findings of monkey area 5, showing that neurons only become activated after the hand of the reach is specified, but not if a target is presented without the hand being specified (Cui & Andersen, 2011). However, it could also be possible that the substantial differences in expected utility between contralateral and ipsilateral arm movements, due to the eccentric location of the target, biased the competition for selection to the contralateral motor plan in Bernier et al.’s study (2012).

Other studies do suggest competition between motor plans of the two hands. Reaction times are longer for reaches towards the target direction that leads to equiprobable right/left hand choices (point of subjective equality, PSE), resembling a more competitive hand selection process for this direction compared to other, lateral target directions (Bakker et al., 2018; Oliveira, Diedrichsen, Verstynen, Duque, & Ivry, 2010). Also, preparing reaches with two hands simultaneously results in more movement variability than preparing a single reach, suggesting that reach plans of the two hands share a common neural resource (Oostwoud Wijdenes, Ivry, & Bays, 2016). Using transcranial magnetic stimulation over left posterior parietal cortex, Oliveira et al. (2010) demonstrated that the competition between hands can be biased towards the ipsilateral, left hand. Fitzpatrick et al. (2019) reported greater BOLD activity in parietal cortex at the PSE than away, consistent with competition between the hands. Finally, using EEG, Hamel-Thibault et al. (2018) presented evidence that hand selection at the PSE depended upon the phase of delta-band oscillations at target onset in contralateral motor regions, as if excitability of motor regions acts as a modulatory factor for hand choice.

Given the importance of beta-band synchronization in movement planning, here we examine the role of these oscillations in coding multiple movement plans during hand choice. Participants performed a hand choice reaching task whereby the target location was cued 1000-1500 ms before it was presented. This allowed us to analyze the oscillatory activity within a clearly defined and motion-artifact-free time window just prior to movement onset. We hypothesized that if beta-band power reflects effector uncertainty, the power would decrease less if there was more uncertainty about which hand to move, similar to the effect of target direction uncertainty (Tzagarakis et al., 2010). We further reasoned that there would be more competition, and thus more uncertainty about which hand to move, if the target was in a direction close to PSE than if the target was close to either of the two hands (Oliveira et al., 2010).

## Methods

### Participants

Twenty participants took part in the study (5 males and 15 females, mean age 21 years, age range 19-26 years). All participants were right-handed, confirmed using the Edinburgh Handedness Inventory (Laterality Quotient, *M* = 86.92, *SD* = 13.54) (Oldfield, 1971). Participants had normal or corrected-to-normal vision and reported no history of neurological or psychiatric diseases, or use of psychoactive medication or substances in the month prior to participation. The ethics committee of the Faculty of Social Sciences of Radboud University Nijmegen, the Netherlands, approved the study. All participants gave written informed consent prior to the start of the study.

### Setup

Participants were seated in front of a touch screen, positioned in the horizontal plane at the level of their thoracic diaphragm. The screen had a resolution of 1920 x 1080 pixels (pixel pitch 0.4845 mm) and a refresh rate of 60 Hz. As illustrated in Figure 1A, two starting positions for the left and right index finger were presented as gray discs of 3.5 cm diameter, approximately 20 cm away from the participant’s sternum and 9 cm on either side of the body midline. A white fixation cross with a width of 2.5 cm was presented along the body midline, 12 cm in front of the two start positions. Cues and targets were presented as light orange and blue 3.5 cm discs, respectively, at 30 cm distance from the point midway between the two start positions, in five different directions: −40°, −10°, 0°, 10°, 40°. A 64-channel active electrode EEG system was used to record brain activity (Brain Products, Gilching, Germany). The onset of visual stimuli on the touch screen was determined using a photodiode and was used to identify and align epochs in the EEG recording. Horizontal and vertical electro-oculograms (EOGs) were recorded by placing electrodes at the supraorbital and infraorbital ridges of the left eye and the outer canthi of the left and right eye. Impedance values for all electrodes were kept below 20 kΩ and the signal was referenced against the signal on left mastoid electrode TP9. The data were filtered online with a low cutoff value of 0.016 Hz and a high cutoff value of 200 Hz and digitized with a sampling frequency of 500 Hz and a resolution of 0.1 μV. The experiment was controlled using custom-written software in Python.

**Figure 1.**
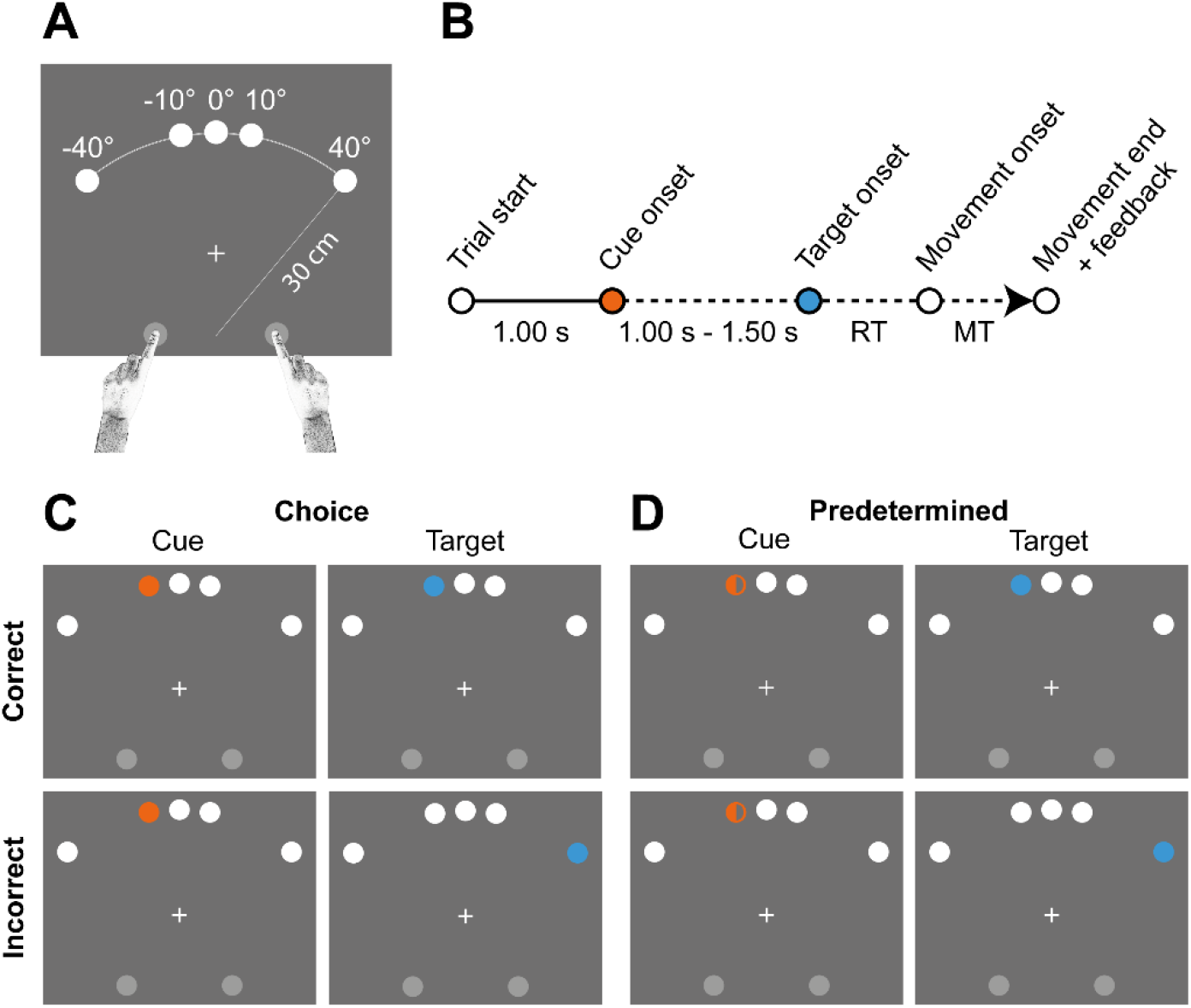
Illustration of the experimental set-up, procedure, and paradigm. A) Schematic illustration of the experimental set-up. Start positions (gray disks), gaze fixation cross, and the five potential cue and target directions (white disks) are shown. B) Order of events in a single trial. C) Choice trials; the upper panels show a correctly cued trial, during which the cue (orange) appeared at the same position as the target (blue), the lower panels show an incorrectly cued trial, during which the target appeared at a different position than the cue. Note that the other potential cue and target directions were not shown during the experiment. D) Predetermined trials; same as in C), but here the cue stimulus instructed which hand to use (here: left hand).

### Paradigm

The experiment took place in a completely darkened room, except for the light of the touch screen. Participants performed a unimanual reaching task in which they were free to use either hand (choice trials) or in which the response hand was instructed on the screen (predetermined trials). All trials were initiated by asking participants to place the tips of their left and right index fingers on the starting positions, which then turned white, and look at the fixation cross. After a delay of 1 s one of the five target directions was cued for either 1.00, 1.25, or 1.50 s (Fig 1B). Presented as a full orange disk, the cue instructed a choice trial (Fig 1C); if the color filled half of the disc, it signaled a predetermined trial (Fig 1D), with the filled side (left or right) instructing which hand to use. Participants were informed about the types of cues prior to the experiment and practiced this before the start of the experiment. Furthermore, the cue was either valid in terms of the upcoming target direction (i.e., correctly cued the target, Fig 1C and 1D, upper panels) or invalid (Fig 1C and 1D, lower panels). At target presentation the cue disappeared and a short beep was played. Participants were asked to touch the target as fast as possible while the eyes were free to move. To ensure that participants were motivated to reach toward the target quickly, they received a feedback message and a score after each response. If participants adequately touched the target within 0.7 s (i.e., reaction + movement time) the message read, ‘Well done! +1 point’, followed by the total earned score across trials. If this duration was beyond 0.7 s, the feedback message was ‘Too slow’, and no points were obtained. If the movement was initiated prior to the onset of the target, the trial was restarted. The incorrectly cued trials serve to verify that motor planning occurred during the cueing phase rather than participants waiting for the target to start preparing their movement.

Each participant completed 900 trials in total, which took about one hour. These comprised of 450 correctly cued trials (90 repetitions of each of the five locations) and 450 incorrectly cued trials (22 or 23 repetitions of each of the 20 cue x target combinations). There were 800 choice trials and 100 predetermined trials, of which 50 left hand and 50 right hand trials (25 correctly cued trials and 25 incorrectly cued trials each). For each participant, trials were presented in a random order in six blocks of 150 trials, separated by short breaks. Prior to the main experiment, participants performed 30 practice trials, including all trial types.

### Data analysis

#### Behavioral analysis

Behavioral data were processed in MATLAB R2017a. Statistical analyses were done in R 4.0.1 and the alpha level was set to 0.05. Choice data were based on the touch screen measurements. Movement onset was defined as the moment the first hand released contact with the touch screen after the target was presented. Hand choice was determined as the hand that departed first. Trials during which the participant released both hands and predetermined trials during which the participant did not use the instructed hand were not taken into account in further analyses. On average, this was the case in 8 trials per participant (*SD* = 3.22). Hand choice preferences were quantified as the proportion of right hand choices for each target direction.

Although there were only five cue and target directions, we summarized the psychometric data for the correctly cued choice trials by fitting a cumulative Gaussian distribution per participant using a maximum likelihood approach (Wichmann & Hill, 2001):

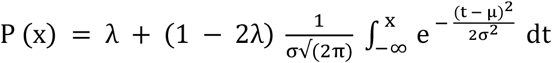

in which P (x) represents the proportion of right hand choices for cue and target direction x. The mean of the curve, μ, represents the participant’s PSE, i.e. the direction at which the right and left hand were chosen equally often. Parameter σ is the standard deviation of the Gaussian, and reflects the variation in choice behavior. Parameter λ represents the lapse rate, accounting for errors caused by participant lapses or mistakes, e.g. unduly reaching with the right hand to the most leftward target. Its value was restricted to small values (< 0.1). We equated the cue direction closest to the PSE direction as the direction that evoked the highest effector competition. Note that the fitted cue direction corresponds to the direction for which the proportion of right hand choices is closest to 0.5 for all participants. Data from three participants were excluded as they showed such a strong preference to reach with their dominant right hand that it was not possible to fit a cumulative Gaussian function, and therefore to select a PSE cue. The extreme left and right directions induced the lowest effector competition. For plotting purposes we also fitted a cumulative Gaussian distribution to the proportion of right hand choices for the five different cue and target directions averaged across participants.

The incorrectly cued choice trials tested whether participants planned movements during the cueing phase. If participants instigated reach planning upon cue presentation, we expect that this would affect the reach upon target presentation. To test if cue direction affected hand choice a cue direction (−40°, −10°, 0°, 10°, 40°) x target direction (−40°, −10°, 0°, 10°, 40°) repeated-measures ANOVA was performed on the proportion of right hand responses for all choice trials (ez package in R). F statistic values were adjusted for violations of sphericity with Greenhouse-Geisser corrections.

Reaction time (RT) was defined as the time between target onset and movement onset. Trials with reaction times <100 ms or >1000 ms were excluded from further analyses. On average, this was the case in 1 trial per participant (*SD* = 1.19). Movement time (MT) was defined as the time between movement onset and the time when the finger first touched the target. Trials with movement times > 1000 ms were excluded from further analyses since these typically involved corrective movements. On average, this was the case in 9 trials per participant (*SD* = 17.92). To test if effector competition was reflected in reaction times, a linear mixed-effects model with participant number as a random factor with random intercept and fixed factors instruction (predetermined, choice), cue direction (PSE, extreme), cue validity (correct, incorrect), and cue time (1.00, 1.25, 1.50 s), as well as the interaction effects, was fitted to the reaction times of all trials using maximum likelihood estimation (nlme package in R). Model fits were assessed with a likelihood ratio test. Bonferroni corrected pairwise t-tests were used to further analyze significant interaction effects post hoc.

#### EEG analysis

EEG data were processed offline using the MATLAB software toolbox FieldTrip, version 20171130 (Oostenveld, Fries, Maris, & Schoffelen, 2011). Data were split into epochs aligned to the onset of the cue (t = 0 s) and the signal was re-referenced against the average signal of the EEG electrodes. Slow drifts in the signal were eliminated by applying a high-pass filter with a cutoff frequency of 1 Hz. Eye blinks were semi-automatically identified based on the difference signal between the two vertical EOG electrodes following the FieldTrip procedure for rejection of eye blink artifacts. Trials with eye blinks around the onset of the cue (time window from 75 ms prior to cue onset to 25 ms after cue onset) were removed from further analyses. On average, this resulted in removal of 18 trials per participant (*SD* = 16.89). Ocular artifacts during the remainder of the trial were removed from the signal by running an independent component analysis. Rejection of components with an evident ocular origin was done according to the criteria described by McMenamin et al. (2010). After removal of these components, trials with excessive muscle activity in the time window from 200 ms prior to cue onset until target onset were semi-automatically identified and removed from further analyses following the FieldTrip procedure for rejection of muscle artifacts (see Gonzalez-Moreno et al., 2014 for further details). On average, this resulted in removal of 102 trials per participant (*SD* = 50.76). Bad channels were identified by visually inspecting the preprocessed data and were repaired by replacing the data with the plain average signal of neighboring channels based on triangulation (two channels repaired in total). Data were low-pass filtered with a cutoff frequency of 40 Hz and down-sampled to 200 Hz.

Time-frequency representations of the data were computed with a Hanning taper with variable window length (5 cycles of the frequency of interest per time window), 10 ms steps and a 1 Hz resolution. The procedure was repeated with the epochs realigned to the onset of the movement (t = 0 s). Power values were corrected relative to a baseline computed per participant, trial group, frequency bin and channel. This baseline was defined as the average power in the time window from 200 ms before cue onset until cue onset, and was computed after averaging across trials in a trial group. Baseline-corrected power values were expressed in decibels.

First, we sought to identify clusters of channels that showed activity related to movement preparation. More specifically, we performed a nonparametric cluster-based permutation test to find clusters of channels that showed a decrease in power in the beta-band frequency range (13 to 30 Hz) prior to either left or right hand responses. Trials for which the hand to use was predetermined were grouped based on the hand used (left or right hand). Both correctly and incorrectly cued trials were included, as we did not expect cue direction to affect which hand was prepared for these predetermined trials. We used a nonparametric cluster-based permutation test to find clusters of channels that showed contrasting activity prior to left and right hand movements. This cluster-based permutation test is based on the calculation of cluster-level statistics, connecting samples that are adjacent in space and time (Maris & Oostenveld, 2007). To contrast left and right hand trials, power values in the right hand trial group were subtracted from the power values in the left hand trial group. The remainder was averaged along the frequency dimension within the beta-band range (13 to 30 Hz). The permutation test was applied for the channels in the left and right hemisphere separately, and channels were spatially clustered using triangulation of the sensor positions. Clusters in time were restricted to occur in the time window from 500 ms before movement onset until movement onset, mainly overlapping with the reaction time window. Both the cluster alpha level and the alpha level to reject the null hypothesis of no clusters in the data were set to 0.05. Mirror-symmetric channels that could be found in a significant cluster in the left hemisphere as well as a significant cluster in the right hemisphere were selected for further analyses, and data were averaged across the channels within a channel cluster.

Second, we were interested in whether effector competition was reflected in beta-band power during motor planning. Trials were grouped based on instruction (predetermined, choice), cue direction (extreme, PSE) and hand used (left, right). For the predetermined trials, both correctly and incorrectly cued trials were included. For the choice trials, only the correctly cued trials were included, as participants might have chosen to switch hands after the presentation of an incorrectly cued target, making it inappropriate to group trials based on the hand used. For reaches towards the extreme cues, only left hand trials were included for the leftmost cue (−40°) and only right hand trials were included for the rightmost cue (40°). Power values were computed for the sensor clusters ipsilateral and contralateral to the hand used, and were collapsed across hands, resulting in trial groups based on instruction (predetermined, choice), cue direction (extreme, PSE) and sensor cluster (contralateral, ipsilateral). Power values were averaged along the frequency dimension in the beta-band range (13 to 30 Hz).

To test if beta-band power was modulated by instruction, cue direction and sensor cluster, we performed a repeated-measures ANOVA on the average beta-band power during the time window from cue onset until 1000 ms after cue onset, with instruction (predetermined, choice), cue location (extreme, PSE), and sensor cluster (ipsilateral, contralateral) as factors. A Bayesian ANOVA was used to compute Bayes factors for all main and interaction effects (BayesFactor package in R, see also Rouder, Morey, Speckman, & Province, 2012). To examine whether the effects were limited to the power in the beta-band frequency range, the procedure was repeated for the power in the theta-band (5 to 7 Hz) and alpha-band frequency range (8 to 12 Hz).

## Results

To examine if cortical power reflects uncertainty in hand choice, participants performed a cued hand choice reaching experiment, whereby the hand to use was chosen by the participant, based on a cue and target, or instructed by the cue. Figure 2A shows the proportion of right hand choices for the five different target directions when correctly cued averaged across participants (open circles) and their psychometric fit (in black), superimposed on the fits of individual participants with their PSEs (gray circles). Confirming previous literature (Bryden, Pryde, & Roy, 2000; Gabbard & Rabb, 2000), the ipsilateral hand was typically selected to reach for peripheral targets, i.e. the left hand reached to the - 40° target, the right hand reached to the 40° target. Most participants had a negative PSE, indicating an overall bias to selecting the right hand, which is consistent with the right hand preference of our participants. The direction closest to the participants’ PSE was selected as the high competition direction: −10° (n = 13), 0° (n = 3), or 10° (n = 1). We will refer to this direction as the participant’s *PSE cue or target*.

**Figure 2.**
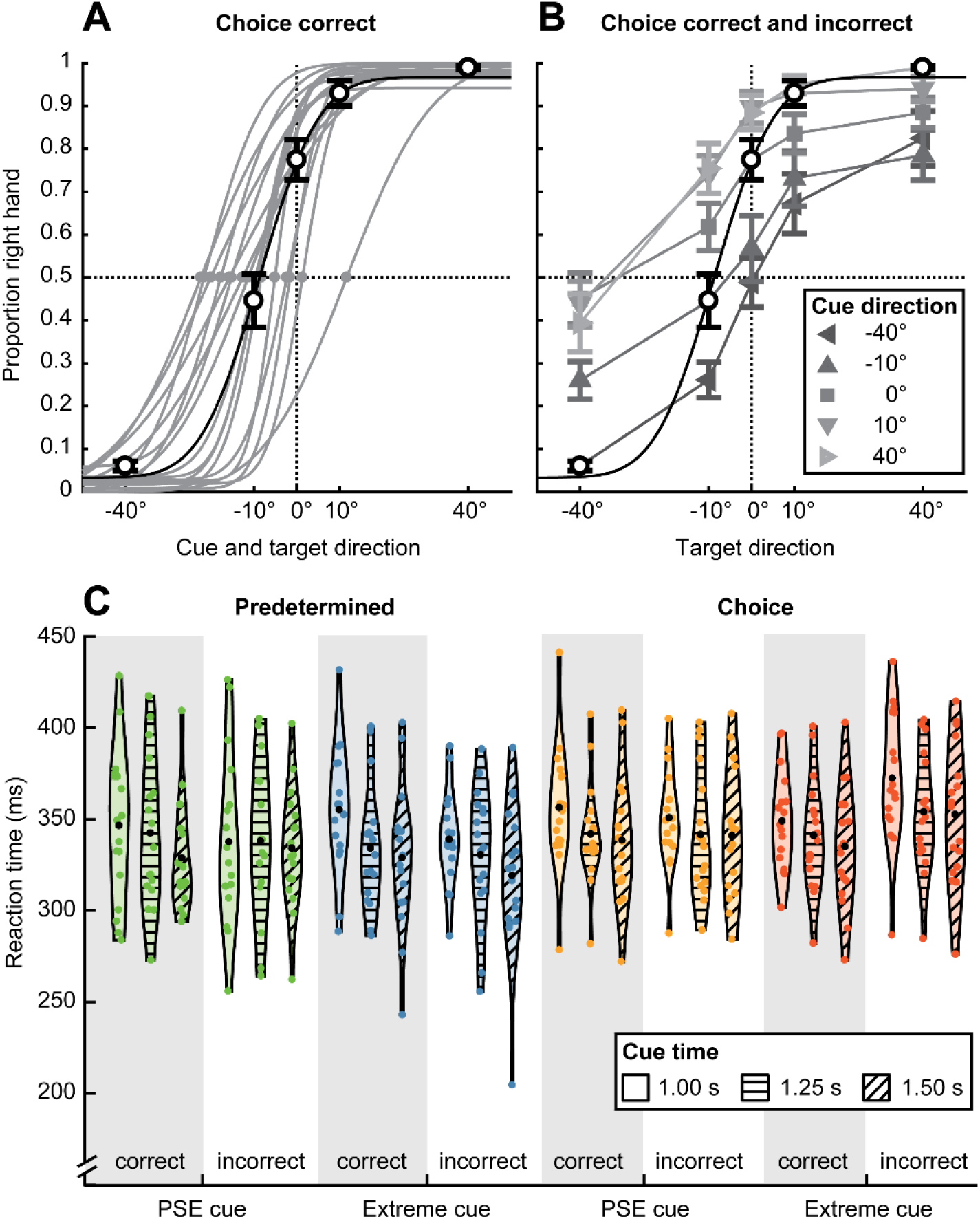
Choice behavior and reaction times. (A) Proportion of right hand choices as a function of cue and target direction for correctly cued choice trials (open circles) fitted with a cumulative Gaussian distribution for all participants (black line). Points of subjective equality (gray dots) and cumulative Gaussian fits for individual participants (gray lines). (B) Proportion of right hand choices as a function of cue (gray lines) and target direction (abscissa) for correctly (open circles; same as in panel A) and incorrectly cued choice trials (triangle and square shapes) for all participants. A repeated-measures ANOVA was used to examine the effect of cue and target direction on hand choice (n = 17). (C) Reaction times as a function of instruction, cue direction, cue validity and cue time for all participants. Violin shape outlines show the kernel density estimates of the individual participant data points (colored dots). Black dots show the mean across participants. A linear-mixed effects model was used to examine the effect of instruction, cue direction, cue validity and cue time on reaction times (n = 17).

We used the incorrectly cued choice trials to find behavioral evidence for motor planning during the cueing phase. We reasoned that if participants simply postponed motor planning until the presentation of the target, the cueing phase should not affect response behavior. Alternatively, if motor planning occurs in the cueing phase, it should bias hand choice. Figure 2B shows that cue location affects hand choice. For example, for a −40° cue and target (leftmost open circle), participants almost invariably use the left hand, while for the −40° target in combination with other cue locations the subsequent hand choice is more ambiguous. Similar effects can be seen across all invalid cue-target combinations. Thus motor planning during the cueing phase affected later hand choice. In support, across all choice trials, a repeated-measures ANOVA showed significant main effects of cue (*F*(1.68, 26.83) = 27.02, *p* < 0.001) and target direction (*F*(1.66, 26.57) = 102.18, *p* < 0.001) on hand choice, as well as a significant interaction (*F*(7.04, 112.60) = 7.60, *p* < 0.001). This confirms that the cue affects the eventual response, justifying our choice to study movement preparation during the cue period.

To test whether the paradigm evokes competitive processes in which both hands compete for movement execution we performed a reaction time analysis. Figure 2C shows the reaction times for the different conditions. A linear mixed-effects model fitted on the reaction times with fixed effects instruction (predetermined, choice), cue direction (PSE, extreme), cue validity (correct, incorrect) and cue time (1.00, 1.25, 1.50 s) showed a main effect of instruction, illustrating longer reaction times for choice trials than for predetermined trials (*χ^2^*(1) = 31.56, *p* < 0.0001). This suggests that, as expected, there was more competition between the hands for choice trials than for predetermined trials. There was also a main effect of cue time (*χ^2^*(2) = 45.02, *p* < 0.0001). Post hoc tests revealed that reaction times were longest for the shortest cue period (*M* = 349 ms) and shortest for the longest cue period (*M* = 332 ms) (*p* < 0.0001), suggesting that motor preparation might have been further advanced with longer cue times, leading to shorter reaction times.

Based on Oliveira et al. (2010) we hypothesized that reaction times would be longer for the PSE cue than for the extreme cues, but there was no main effect of cue direction on reaction time. However, there were two significant interaction effects with the factor cue direction: the two-way interaction between instruction and cue direction (*χ^2^*(1) = 5.37, *p* = 0.021) and the three-way interaction between instruction, cue direction and cue validity (*χ^2^*(1) = 12.20, *p* < 0.001). The two-way interaction seems to be driven by longer reaction times for choice trials than predetermined trials if the cue was in an extreme direction (*p* = 0.16), rather than if the cue was in the PSE direction (*p* = 0.66). The three-way interaction suggests that this effect was driven by the incorrectly cued trials. Overall, reaction times were not longer for the PSE cue than for the extreme cues. However, for incorrectly cued choice trials, reaction times were longer for the extreme cues than for the PSE cue. This suggests that there might not have been more competition between the hands, and thus more uncertainty about hand choice, for PSE than for extreme cues.

Finally, there was a significant interaction effect of instruction and cue validity on reaction time (*χ^2^*(1) = 15.36, *p* < 0.0001), demonstrating that incorrect cues only prolonged reaction times for choice trials (*p* < 0.0001), but not for predetermined trials (*p* = 0.064). Most likely participants did switch hands from cue to target in choice trials, while switching was not allowed in predetermined trials. This is further support for participants not postponing motor planning until the presentation of the target.

We next turned to examining the cortical mechanisms, studying whether power changes in motor planning regions reflect uncertainty about the upcoming effector. Our focus is on the role of beta-band oscillations, known to be involved in motor planning, and implicated in the coding of multiple target-specific motor plans. We used the predetermined trials to select the cortical regions that show beta-band activity during left and right hand motor planning around movement onset. As shown in Figure 3, we found two clusters of sensors that showed a significant selectivity in the beta band for the contralateral hand, one in the left (*p* = 0.039) and one in the right (*p* = 0.021) hemisphere. The mirror-symmetric channels that could be found in both significant clusters mostly covered central areas of the brain. Across the left hemisphere these channels were FC1, C1, C3, C5, T7, CP1 and CP5, and across the right hemisphere these channels were FC2, C2, C4, C6, T8, CP2 and CP6. These clusters are centered around central channels C3 and C4, known to be involved in movement planning (Pfurtscheller, 1992).

**Figure 3.**
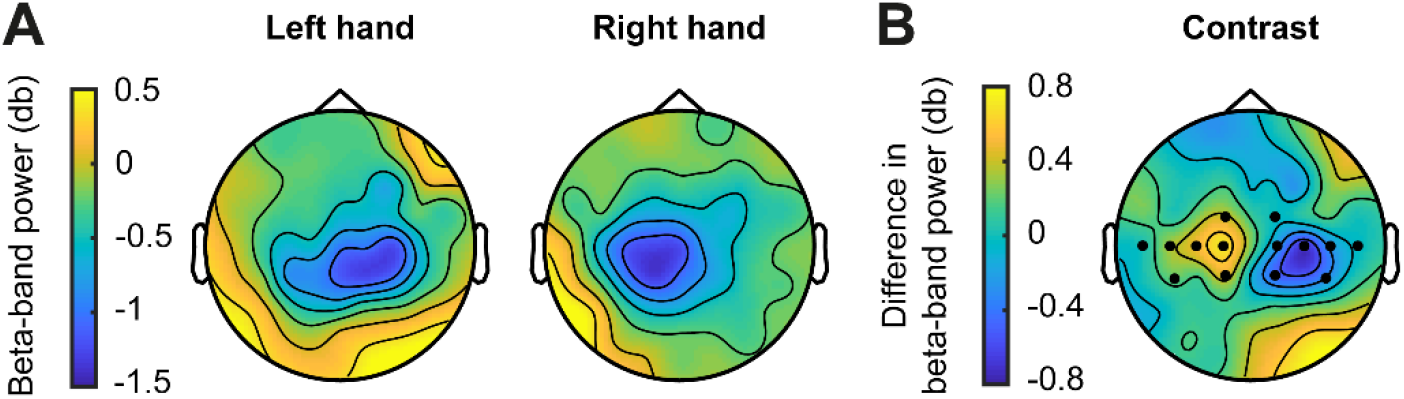
Topographic map of beta-band power preceding left and right hand movements. (A) Mean beta-band power for the predetermined trials (correctly and incorrectly cued) preceding left and right hand movements (time-locked to movement onset, averaged across the 500 ms preceding movement onset). (B) Mean difference in beta-band power between the hands (left minus right hand). Channel clusters (black dots) were identified with a nonparametric cluster-based permutation test (n = 17).

We next examined whether beta-band power during the cueing phase reflects a hand selection process. Figure 4 illustrates relative beta-band power as a function of time, aligned to cue onset (left panels) and response onset (right panels), for both the choice and predetermined trials at the PSE and extreme cues, separately for sensor clusters ipsilateral and contralateral to the selected hand. While there appears a clear difference after cue presentation between choice and predetermined trials in the contralateral cluster, this effect is less pronounced in the ipsilateral cluster. In the contralateral cluster, the power in the beta-band after onset of the cue decreased more in predetermined than choice trials; this difference is sustained until response onset, and appears slightly larger for cues at PSE than at an extreme location. An instruction (predetermined, choice) x cue direction (PSE, extreme) x sensor cluster (ipsilateral, contralateral) repeated measures ANOVA revealed significant main effects of sensor cluster (*F*(1, 16) = 40.61, *p* < 0.0001), consistent with the contralateral selectivity, and instruction (*F*(1, 16) = 20.14, *p* < 0.001), consistent with a smaller decrease in beta-band power in choice trials than in predetermined trials. This suggests that a larger decrease in beta-band power corresponds to less effector uncertainty.

**Figure 4.**
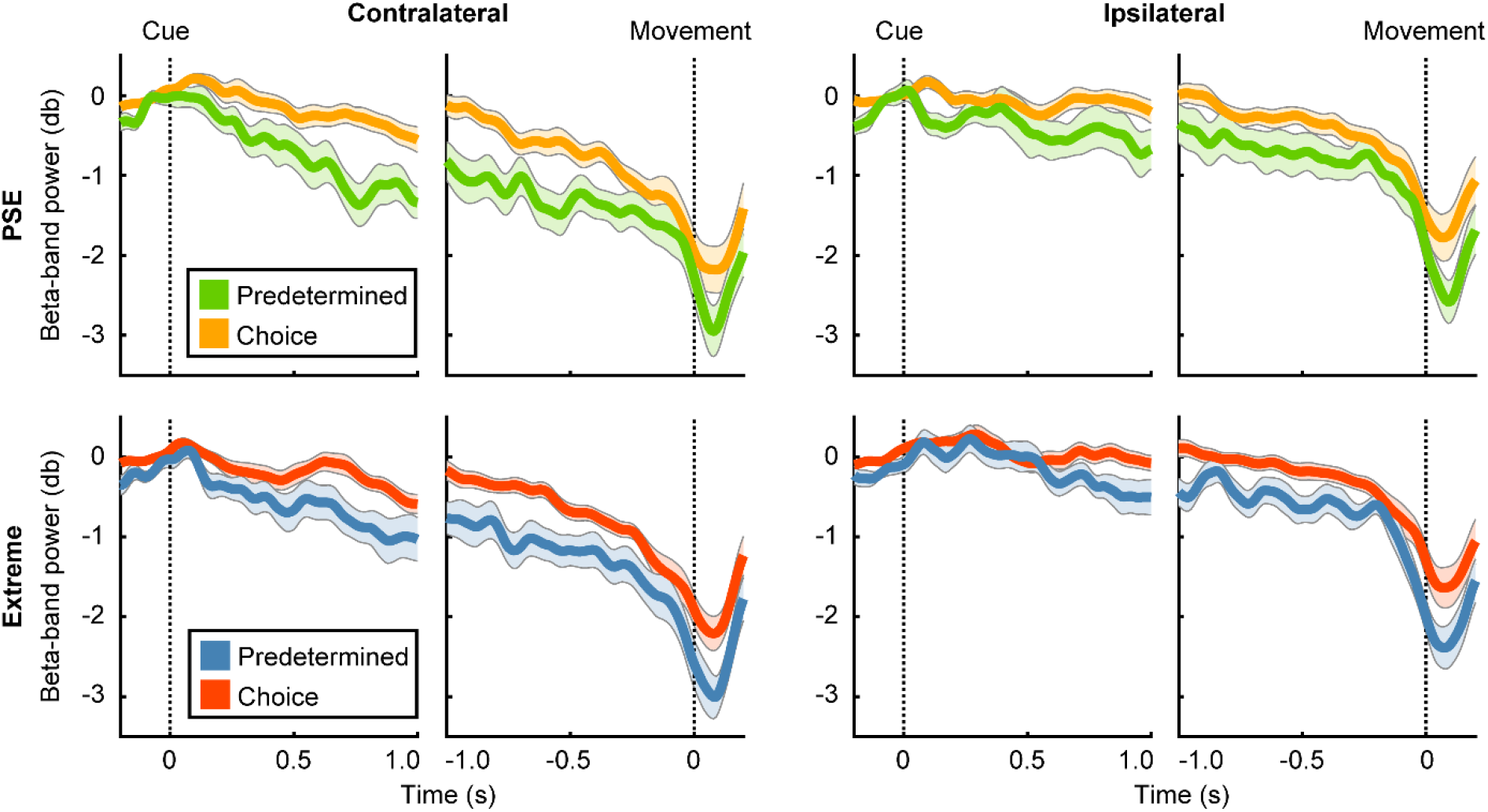
Beta-band power. Relative beta-band power as a function of time in the contralateral (left columns) and ipsilateral (right columns) sensor cluster for the PSE (upper row) and the extreme cue (bottom row). Left and right subpanels show the signal aligned to cue and movement onset, respectively. Shaded areas represent SEM. A repeated-measures ANOVA with the average beta-band power during the time window from cue onset until 1 s after cue onset was used to examine the effect of instruction, cue location and sensor cluster (n = 17).

There was no main effect of cue direction on beta-band power (*F*(1, 16) = 2.30, *p* = 0.149) nor were there any significant interaction effects. One could expect that in the contralateral hemisphere, for choice trials but not predetermined trials, there would be more competition between the hands, and thus more uncertainty, for PSE than for extreme cues. However, an instruction (choice, predetermined) x cue direction (PSE, extreme) repeated measures ANOVA on beta-band modulation in the contralateral cluster only showed a significant main effect of instruction (*F*(1, 16) = 29.67, *p* < 0.0001). The interaction between instruction and cue direction was not significant (*F*(1, 16) = 0.64, *p* = 0.436). Also a Bayesian ANOVA revealed a Bayes factor for the interaction between instruction and cue direction of 0.423, which can be interpreted as inconclusive evidence (Jeffreys, 1961). Overall, our results suggest that beta-band power does not only reflect directional uncertainty, as described by Tzagarakis et al. (2010, 2015), but also effector uncertainty induced by hand choice.

To examine whether the effect of effector uncertainty is specific to the signal in the beta-band, we performed the same analysis in the alpha (8 to 12 Hz) and theta-band (5 to 7 Hz) frequency range. Power in the alpha-band is known to show a similar reduction to beta-band power prior to movement onset (Pfurtscheller, 1992). However, alpha-band power does not modulate with directional uncertainty about the upcoming movement (Tzagarakis et al., 2015). Figure 5A shows the power in the alpha-band as a function of time, grouped based on instruction (predetermined, choice), cue direction (extreme, PSE), and sensor cluster (ipsilateral, contralateral). A repeated-measures ANOVA on the average alpha-band power during the cue phase did not reveal any significant main effects of instruction (*F*(1, 16) = 2.77, *p* = 0.116), cue direction (*F*(1, 16) = 0.77, *p* = 0.393), or sensor cluster (*F*(1, 16) = 4.46, *p* = 0.051), or any significant interaction effects.

**Figure 5.**
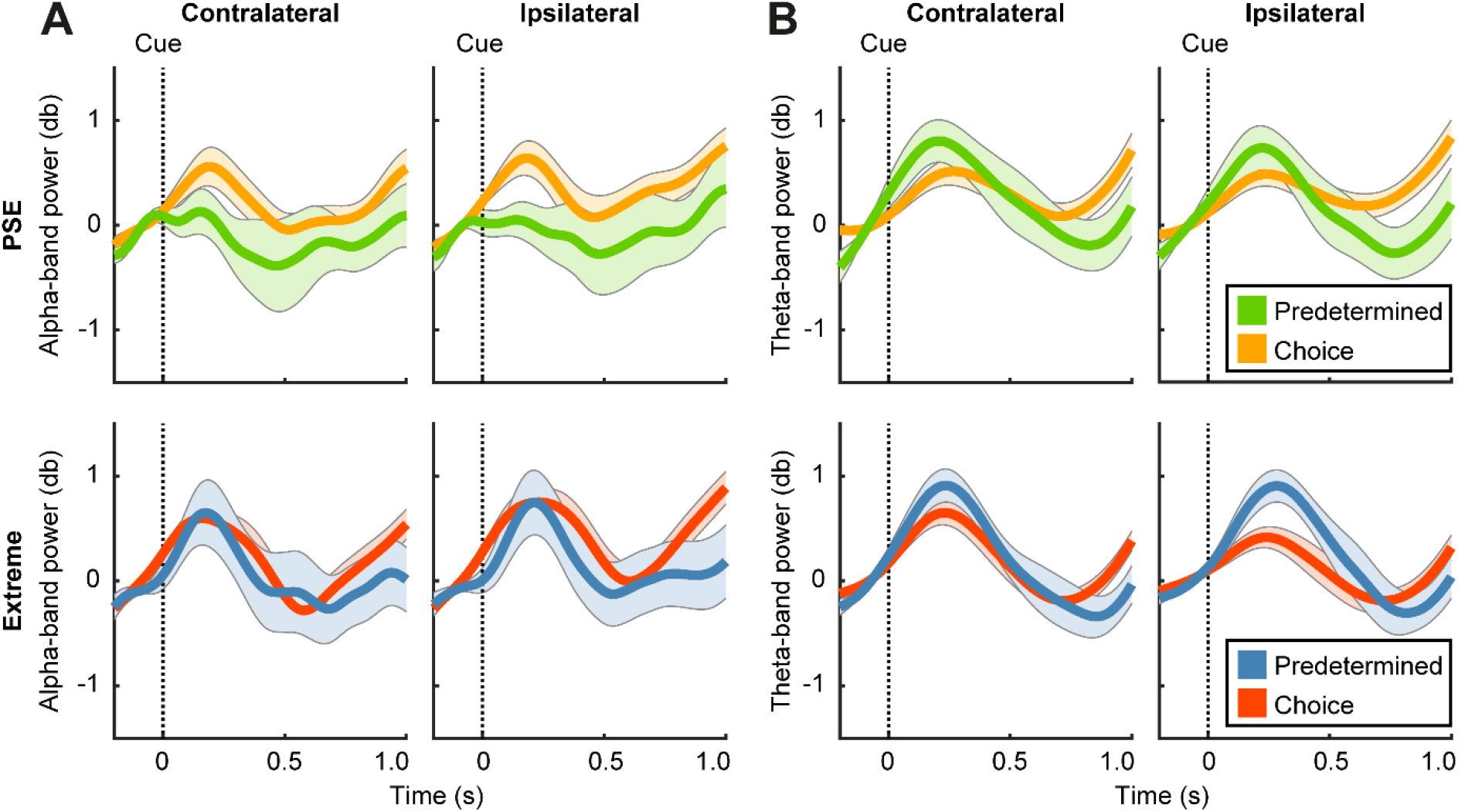
Alpha-band and theta-band power. (A) Relative alpha-band power as a function of time aligned to cue onset in the contralateral (left columns) and ipsilateral (right columns) sensor cluster for the PSE (upper row) and extreme cue (bottom row). Shaded areas represent SEM. (B) Relative theta-band power as a function of time. Configurations the same as panel A. Repeated-measures ANOVAs with the average alpha-band and theta-band power during the time window from cue onset until 1 s after cue onset were used to examine the effect of instruction, cue location and sensor cluster (n = 17).

Next, we examined the effect of effector uncertainty on the oscillations in the theta-band, which have been implicated in motor planning and anticipation (Dufour, Thénault, & Bernier, 2018; Perfetti et al., 2011). Figure 5B shows the power in the theta-band as a function of time during the cueing phase. A repeated-measures ANOVA did not reveal significant main effects of instruction (*F*(1, 16) = 0.00, *p* = 0.967), cue direction (*F*(1, 16) = 0.85, *p* = 0.371), or sensor cluster (*F*(1, 16) = 0.45, *p* = 0.514), or any interactions. The effect of effector uncertainty on oscillations during movement preparation, as recorded from central sensors, thus seems to be specific to the signal in the beta-band frequency range.

## Discussion

To investigate the effect of effector uncertainty on beta-band oscillatory activity during motor preparation, participants performed a hand reaching task whereby the effector to use was either predetermined or free of choice. We hypothesized that competition between the left and right hand would be low, independent of the cue direction, if the hand to be used was predetermined. If participants were free to choose a hand, we expected greater competition and hence a smaller decrease in beta-band power. Additionally we expected more competition during hand choice for the PSE cue than for eccentric cues. Results indicate that effector competition indeed affects beta-band power during motor planning: when participants were free to choose the hand to use beta-band power decreased less than when the hand to use was predetermined. We did not observe a significant effect of cue direction on beta-band power.

Our results demonstrate that effector uncertainty induced by instruction affected beta-band power over central brain areas during motor planning. More specifically, beta-band power decreased less when participants were free to choose the hand to use than when the hand was predetermined. Lower levels of beta-band power are thought to be associated with a readiness to move (Khanna & Carmena, 2017). This idea is in line with our expectations, as the instruction to use a specific hand should diminish competition between left and right-hand motor plans, and therefore ease motor planning. This is further underlined by the observation that instruction also affected reaction times: reaction times were longer when participants were free to choose the hand to use than when the hand was predetermined. This reaction time pattern has been previously observed by Oliveira et al. (2010) and is thought to show that hand selection comes with a cost. All in all, beta-band power was affected by effector uncertainty induced by instruction, with a smaller decrease in power when participants chose the hand for the ensuing reach.

Contrary to our expectations, our results do not show an effect of cue direction, neither on beta-band power, nor on reaction times. We expected that reaches towards the PSE would elicit more competition between the left and right hand than reaches towards targets in the periphery, for which one hand is usually clearly preferred over the other (Bakker et al., 2018; Oliveira et al., 2010; Stoloff, Taylor, Xu, Ridderikhoff, & Ivry, 2011). Indeed, Oliveira et al. (2010) reported that a reaction time difference disappeared by restricting reaches to only one hand, as we do in the predetermined condition. Our experimental paradigm seems to have failed to elicit a difference in effector competition for the PSE and extreme targets. A potential reason can be found in the introduction of incorrect cues, which could have unintendedly increased uncertainty about the effector to use. For every presented cue, there was only 50% chance that the target would be presented in the same direction. Possibly, this resulted in too much uncertainty about which hand to use and therefore participants did not yet fully commit to preparing a single hand. This effect of cue validity on effector uncertainty should be limited to the choice trials, as competition is thought to be low for reaches with a predetermined hand, regardless of the direction of the cue and target. Indeed, our results show that incorrect cues prolong reaction times for choice trials, but not for predetermined trials. Thus, the introduction of the incorrect cues might have resulted in a lack of a difference in effector uncertainty for the PSE and extreme targets for the choice trials, explaining why no effect of cue direction was observed here.

The absence of an effect of cue direction for the choice trials cannot be explained by an overall lack of movement preparation during the cue period. Not only do our results show that incorrect cues prolong reaction times for choice trials, but hand choice was also biased by the direction of the (incorrect) cue. Both findings suggest that participants prepared the movement based on the cue. This is in line with findings from previous delayed response cueing experiments; Tzagarakis et al. (2010, 2015) found that reaction times were longer if the cue was less informative in terms of the direction of the upcoming target, and Oostwoud Wijdenes et al. (2016) showed that movement variability during a reaching movement was larger if the preceding cue did not specify the hand to use.

In our analysis, the effect of effector uncertainty on brain oscillatory activity over central areas of the brain was limited to the power in the beta-band. Even though oscillations in the alpha-band are known to show a similar decrease in power during motor planning to oscillations in the beta-band (Pfurtscheller, 1992), we did not observe a modulation of alpha-band power based on effector uncertainty. This is in line with findings for directional uncertainty where beta-band but not alpha-band power decreases more if target direction is more certain (Grent-’t-Jong et al., 2014; Tzagarakis et al., 2015). Additionally, Rhodes et al. (2018) found that alpha-band power during a cue period only decreases (followed by an increase) if the direction of the upcoming target is unambiguous, suggesting the activity to be related to movement execution processes rather than motor planning. It thus seems as if alpha-band and beta-band power over central areas of the brain reflect complementary but distinct processes, with alpha-band power being insensitive to uncertainty about the upcoming movement.

Theta-band power is known to increase during motor planning (Perfetti et al., 2011), and has been shown to modulate with the anticipation of visual feedback (Dufour et al., 2018). Here, we did not observe a modulation of theta-band power based on effector uncertainty. Thus, the effect of effector uncertainty on oscillatory power during motor planning seems to be reflected in beta-band power specifically, with the reservation that we did not analyze power changes in the gamma band. Van Der Werf et al. (2010) have reported direction-selective synchronization in the 70 to 90 Hz gamma-frequency band, originating from the medial aspect of the posterior parietal cortex, when planning a reaching movement. Future work should address whether gamma-band synchronization also modulates with hand choice.

How the modulation of beta-band power over central areas of the brain coincides with other changes in neural activity observed during effector selection remains to be answered. Here, we focused on beta-band activity from channels positioned along the central coronal plane of the head, covering central areas of the brain. Localizing the exact neural source of this activity, however, was not one of the main objectives of this study. Previous studies have attempted to find the source of neural activity related to effector uncertainty. Hand choice has, for instance, been shown to be related to the phase of delta-band oscillations at the onset of the reach target in the dorsal premotor cortex and primary motor cortex contralateral to the hand used (Hamel-Thibault et al., 2018). Additionally, BOLD activity appears to be modulated by effector uncertainty in parietal cortex (Fitzpatrick et al., 2019), which is in line with the finding that TMS over the posterior parietal cortex biases hand choice (Oliveira et al., 2010). It remains unknown whether these phenomena, distinct in the type of neural activity and source location, are linked, and for example arise from activity in the same neuronal ensembles, or whether these findings arise from independent processes.

In general, motor decisions are thought to be biased by the expected utility of potential movements. This utility depends on the costs and benefits of a certain movement and is based on the location of the movement target relative to the effector. However, also other factors might be taken into account, such as the task or trial instruction. Neural activity related to motor decision making based on utility is thought to intertwine with the activity related to motor planning (Cisek, 2006). Evidence for this has been found in both human (Grent-’t-Jong et al., 2014, 2015; Tzagarakis et al., 2010, 2015) and non-human primates (Basso & Wurtz, 1997; Cisek & Kalaska, 2005; Glaser et al., 2018; Klaes et al., 2011). In line with this, we observe an effect of motor decision making on beta-band power - a neural marker of motor planning (Jasper & Penfield, 1949; Pfurtscheller, 1992).

Our results support the idea that motor plans for the two arms are prepared in parallel and compete for execution, but this idea has been a topic of debate. Bernier et al. (2012) suggested that effector selection actually precedes motor planning. In their experiment, they found activity in the parietal and premotor cortex contralateral to the hand used, but this was only observed after target onset, and thus after the hand was thought to be selected. However, their hand choice experiment differed from the paradigm used here. Bernier et al. (2012) asked participants to reach to two eccentric targets. Additionally, participants never actually chose the hand to use themselves, but were either instructed early on in the trial (based on the cue) or at target onset. Both the location of the targets and the instruction of the hand might have diminished possible competition between left and right hand movement plans, similar as to the predetermined reaches towards an extreme target direction here. It is important to point out though that Bernier et al.’s (2012) findings are in line with results from monkey studies that show that neuronal activity only encodes selected reach plans, instead of potential reach plans, in area 5 (Cui & Andersen, 2011) and dorsal premotor cortex (Dekleva et al., 2018). Based on these results, Dekleva et al. (2018) challenge the idea of the parallel specification of motor plans for potential reaching actions (Cisek & Kalaska, 2005), and suggest that evidence for the encoding of multiple motor plans is simply a result of trial averaging. Unfortunately, we lack the signal-to-noise ratio to address this issue at the single trial level, but this would be an interesting issue for further research.

It could be asked whether the unbalanced number of trials in the predetermined and choice conditions biased our conclusions. While participants completed 100 predetermined trials versus 800 choice trials, we do not believe that participants perceived the predetermined cue stimulus as a deviant. The effect of instruction on beta-band power did not show up just shortly after the presentation of the cue, which might reflect the processing of a surprising visual stimulus, but appeared to be sustained and to even increase throughout the cue period. In support, although the data for the predetermined trials had slightly larger variability than the data for the choice trials, the main effect of instruction on beta-band power was highly significant (*p* < 0.001).

In conclusion, the results of this study suggest that effector competition during motor planning is reflected in beta-band, but not alpha or theta-band, power over central regions. More specifically, beta-band power decreased less with more competition between the left and right hand. Alpha and theta band power lacked these modulations. Our findings support the more general idea that the brain specifies multiple possible effector-specific actions in parallel up to the level of motor preparation.

## Grants

This work was supported by an internal grant from the Donders Centre for Cognition. L.G. was supported by a Veni grant [451-16-013] from the Netherlands Organization for Scientific Research.

## Disclosures

No conflicts of interest, financial or otherwise, are declared by the authors.

## Author contributions

M.J.L.H., L.O.W., L.G. and W.P.M. conceived and designed research; M.J.L.H. performed experiments; M.J.L.H. and L.G. analyzed data; M.J.L.H., L.O.W., L.G. and W.P.M. interpreted results of experiments; M.J.L.H. prepared figures; M.J.L.H., L.O.W. and W.P.M. drafted manuscript; M.J.L.H., L.O.W., L.G. and W.P.M. edited and revised manuscript; M.J.L.H., L.O.W., L.G. and W.P.M. approved final version of manuscript.

## Data availability statement

Upon publication, all data and code will be made publicly available via the persistent identifier currently reserved for this collection: https://doi.org/10.34973/qjgg-h917.

## Notes

https://doi.org/10.34973/qjgg-h917

### Competing Interest Statement

The authors have declared no competing interest.

